# Nucleic Quasi-Primes: Identification of the Shortest Unique Oligonucleotide Sequences in a Species

**DOI:** 10.1101/2023.12.12.571240

**Authors:** Ioannis Mouratidis, Maxwell A. Konnaris, Nikol Chantzi, Candace S.Y. Chan, Austin Montgomery, Fotis A. Baltoumas, Michail Patsakis, Manvita Mareboina, Georgios A. Pavlopoulos, Dionysios V. Chartoumpekis, Ilias Georgakopoulos-Soares

## Abstract

Despite the exponential increase in sequencing information driven by massively parallel DNA sequencing technologies, universal and succinct genomic fingerprints for each organism are still missing. Identifying the shortest species-specific nucleic sequences offers insights into species evolution and holds potential practical applications in agriculture, wildlife conservation, and healthcare. We propose a new method for sequence analysis termed nucleic “quasi-primes”, the shortest occurring sequences in each of 45,785 organismal reference genomes, present in one genome and absent from every other examined genome. In the human genome, we find that the genomic loci of nucleic quasi-primes are most enriched for genes associated with brain development and cognitive function. In a single-cell case study focusing on the human primary motor cortex, nucleic quasi-prime genes account for a significantly larger proportion of the variation based on average gene expression. Non-neuronal cell-types, including astrocytes, endothelial cells, microglia perivascular-macrophages, oligodendrocytes, and vascular and leptomeningeal cells, exhibited significant activation of quasi-prime containing gene associations related to cancer, while simultaneously suppressing quasi-prime containing genes were associated with cognitive, mental, and developmental disorders. We also show that human disease-causing variants, eQTLs, mQTLs and sQTLs are 4.43-fold, 4.34-fold, 4.29-fold and 4.21-fold enriched at human quasi-prime loci, respectively. These findings indicate that nucleic quasi-primes are genomic loci linked to the evolution of species-specific traits and in humans they provide insights in the development of cognitive traits and human diseases, including neurodevelopmental disorders.

## Introduction

Over the last two decades, the cost of sequencing the complete genome of an organism has rapidly declined. Advances in parallel DNA sequencing technologies have enabled the generation of reference genomes for thousands of biological organisms, across viral, archaeal, bacterial, and eukaryotic species (Schoch et al. 2020; O’Leary et al. 2016). The availability of multiple, diverse organismal genomes has enabled advances in bioinformatic analyses and accelerated scientific discovery. Subsequently, our understanding of evolution, encompassing mechanisms of horizontal and vertical gene transfer, selection pressures, and emergence of new species traits, has improved. This information has been utilized in various domains such as human health, pathogen surveillance, agriculture, and species conservation, leading to the development of approaches for disease diagnosis, enhancement of food safety, and mitigation of antibiotic resistance, among other applications (Jagadeesan et al. 2019; Deurenberg et al. 2017; Brandies et al. 2019; Maljkovic Berry et al. 2020). The number of reference organismal genomes is expected to continue to increase and encompass a significant proportion of the genetic diversity present in nature (Darwin Tree of Life Project Consortium 2022; Lewin et al. 2018). For example, the Earth BioGenome Project aims to sequence the genomes of all eukaryotic species within the next ten years (Lewin et al. 2022). This is essential for understanding evolutionary history, discovering the ecological interactions of living organisms and developing precise techniques to detect genomic differences between species. Nevertheless, the increase in biological data also necessitates algorithmic advances to capture the most useful information.

As species evolve and diverge, they acquire new traits. This leads to the divergence of lineage-specific DNA at varying rates as subsets of the genome evolve more rapidly (Seehausen et al. 2014). As an example, after the divergence of humans from other primates, there was an expansion of cranial capacity, brain size and cognitive abilities in humans (Florio, Borrell, and Huttner 2017). Comparative genomics and phylogeny analyses identified regions of organismal genomes that show patterns of accelerated evolution (Foley et al. 2023; Ferris et al. 2018). The usage of mutation rate patterns, species sequence alignments, and the identification of highly conserved regions can provide insights into phenotypic changes (McLean et al. 2011; Hubisz and Pollard 2014). However, available methods rely on genome alignments and mutational analysis, often lacking the ability to identify the units of accelerated evolution at base-pair resolution. The identification of the shortest genomic sequences unique to a species can improve our understanding of how genomic regions evolve and serve as an alternative method for the identification of genomic loci that are changing, at a base-pair resolution.

Kmer analysis involves counting and comparing substrings of length *k* in biological sequences. The distribution of nucleotide kmers varies substantially across organismal genomes (Chor et al. 2009; Bussi, Kapon, and Reich 2021). The presence of individual nucleotide kmers in a species is dependent on several factors, including the GC content of its genome, the biological roles associated with each particular kmer, and the mutation patterns associated with that organism (Bussi, Kapon, and Reich 2021). The set of kmers in a species’s genome can serve as a signature of its underlying sequence. Comparing these kmers and their frequencies enables the identification of distinct characteristics among species. For example, extremophile genomes exhibit some of the most distinct and unique kmer profiles (Bize et al. 2021).

Nullomers are the subset of nucleotide kmers that are not observed in a genome (Hampikian and Andersen 2007), and their absence has been previously attributed to mutational patterns and negative selection (Georgakopoulos-Soares, Yizhar-Barnea, et al. 2021; Koulouras and Frith 2021). Additionally, genome primes are the subset of nullomers that are absent from the genomes of all species (Hampikian and Andersen 2007). Applications have included the usage of nullomers in barcoding (Goswami et al. 2013), deriving anticancer peptides (Alileche et al. 2012), developing vaccine adjuvants (Patel et al. 2012), and in cancer detection (Georgakopoulos-Soares, Barnea, et al. 2021; Montgomery et al. 2023). Therefore, the development of efficient algorithms and methodologies that derive highly informative kmers can result in multiple practical applications.

Previously, we defined a concept termed peptide quasi-primes, the shortest peptide sequences that are unique to a reference proteome (Mouratidis et al. 2023). Here, we extend this concept to nucleic quasi-primes, which are the shortest nucleotide kmers that are unique to a particular species and are nullomers in all other assembled reference genomes (**Figure 1**). We also detect and analyze the set of DNA sequences that are absent from every known genome, also known as DNA primes. As proof of concept, we annotate the set of human quasi-primes and their locations in the human genome. Among *cis*-regulatory elements, quasi-primes are enriched in promoters and 5’UTRs. We observe that human quasi-primes are enriched in genes expressed in the brain and are associated with brain development and neuronal functions. Human quasi-primes are also associated with brain disorders, including schizophrenia, bipolar disorder, intellectual disability, drug abuse, and autism. Finally, we provide evidence that human quasi-primes are enriched for disease variants and variants that significantly impact gene expression, methylation and splicing differences between individuals (**Figure 1**). This work provides the methodology for the identification of nucleic quasi-primes and exemplifies their utility for detecting genomic loci that are associated with human-specific traits and which could be potentially important targets in understanding human brain evolution and brain-associated diseases.

**Figure 1:**
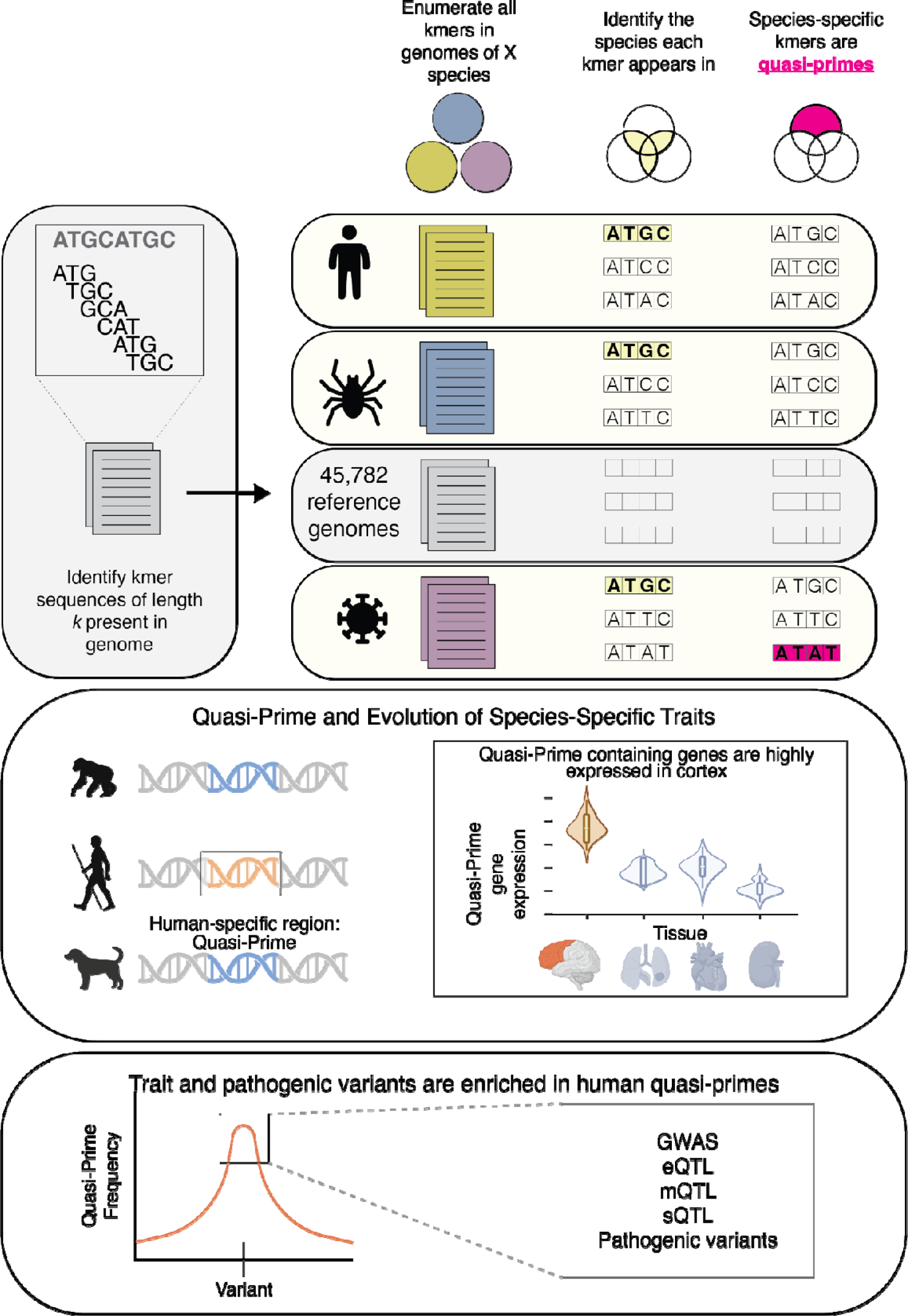
Identification of nucleic quasi-prime sequences in individual species. Schematic displaying the identification of a four-mer quasi-prime sequence in a species. All nucleic kmers of a specific length are identified for each species, across all the reference genomes. Kmers that are shared between multiple species are removed and only kmers appearing in a single reference genome are kept, constituting the set of quasi-primes for that species for that particular length. As an example, we show a short toy sequence that is found in the human reference genome but is absent from all other species in our database, therefore constituting a human nucleic quasi-prime kmer. Quasi-primes are associated with the evolution of human specific traits. For humans, quasi-prime containing genes are enriched in the cortex and are associated with brain development and diseases. Human traits and pathogenic variants are significantly enriched in human quasi-prime sequences. Variants including expression quantitative trait loci (eQTLs), methylation QTL (mQTLs), splicing QTL (sQTLs), genome-wide association studies (GWAS) variants and disease variants are more likely to be found in human quasi-prime sites.

## Results

### Kmer distribution across taxonomic subdivisions and species

We performed a comprehensive analysis of 45,785 reference genomes spanning the three domains of life and viruses. Our aim was to explore the diversity and uniqueness of nucleic sequences across different species and identify kmers that may serve as molecular fingerprints. We first investigated the number of different kmers across each sequenced reference genome as a function of kmer length ranging from one to seventeen bps. For kmer lengths of 16bp, we found that the median number of observed kmers per genome was 3,796,626. We also examined the number of 16bp long kmers detected per genome across taxonomic subdivisions for viral, archaeal, bacterial, and eukaryotic genomes and observed a median of 35,744, 4,912,057, 8,380,257, and 30,028,396 kmers respectively, representing 0.000832%, 0.1189%, 0.1976% and 1.866% of the kmer space (**Supplementary Figure 1**; Figure 2a). These findings were consistent for kmer lengths of fifteen and seventeen bps (**Supplementary Figure 1**; Figure 2a), indicating that the kmer space is sparsely populated by any single reference genome at these kmer lengths. Therefore, we proceeded to investigate the extent to why certain kmers are absent across multiple species and, as an extension, determine if there exist kmers that are unique to a single species.

**Figure 2.**
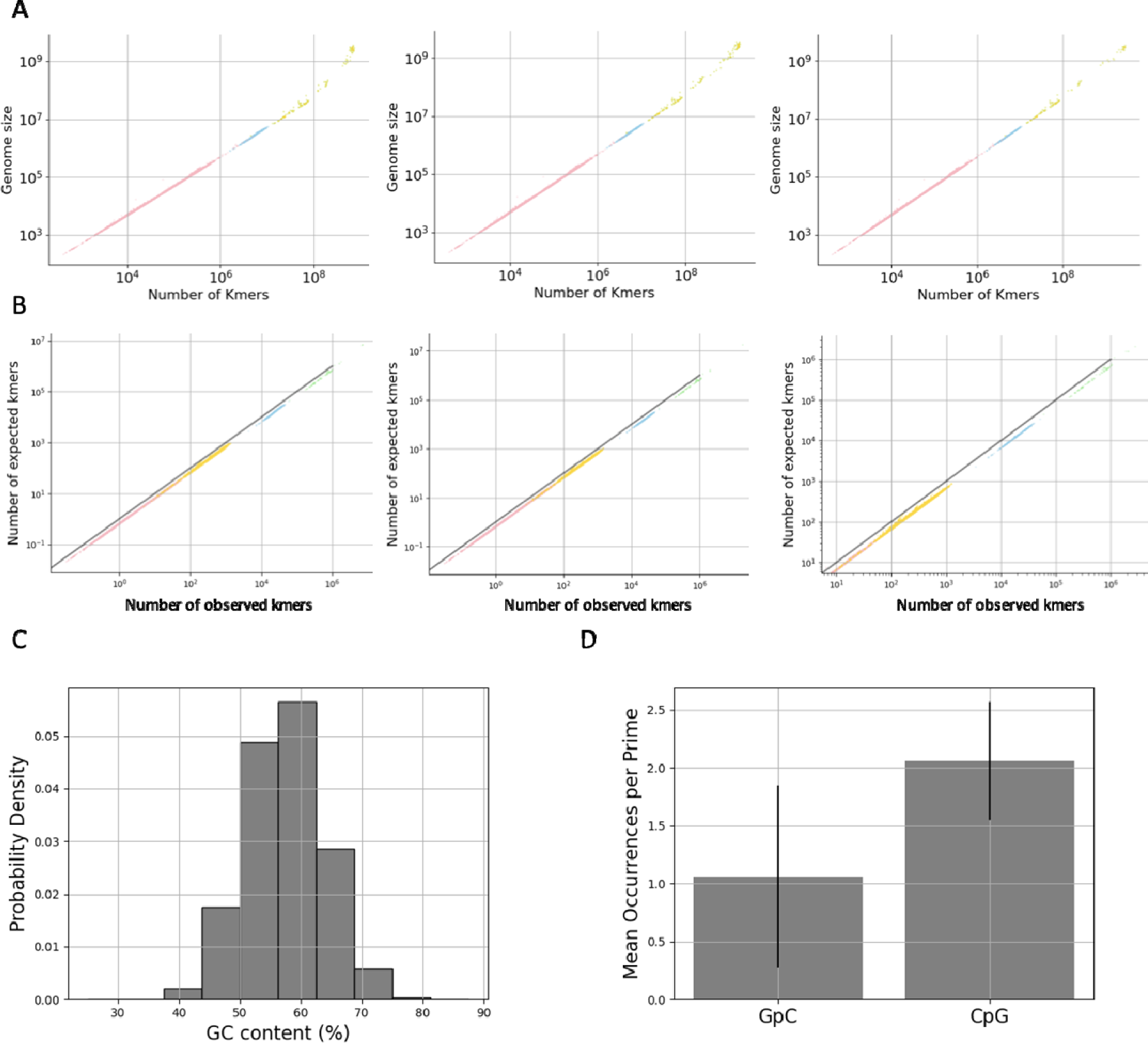
Kmer content as a function of genome length across organisms. **A.** Number of different kmers observed as a function of genome size for kmer lengths of 15bp, 16bp, and 17bp. **B.** Number of expected versus the number of observed kmers for each reference genome for kmer lengths of 15bp, 16bp, and 17bp. Viral, archaeal, bacterial, and eukaryotic genomes are colored pink, blue, yellow, and green respectively across **A-B** figure panels. **C.** GC content percentage of nucleic prime sequences. **D.** Average number of GpC and CpG occurrences per prime. Error bars in D represent standard deviation.

### There are more nullomers than expected across genomes in each taxonomy

In a previous study, we demonstrated that the human genome exhibits a higher prevalence of nullomers than expected by chance (Georgakopoulos-Soares, Yizhar-Barnea, et al. 2021). However, these findings have not been extended to other species, and it is currently unknown if this pattern is consistent across the different organismal genomes present in the taxonomic subdivisions. To address this gap, we performed a comprehensive analysis of the number of expected and observed frequencies of oligonucleotide kmers for each species by generating simulated genomes that account for dinucleotide content (see **Methods**). We then investigated whether there were significant deviations between the two sets of frequencies across different kmer lengths and organisms.

Our results reveal a consistent pattern of nullomer enrichment across species and taxonomic groups. Specifically, for kmer lengths of 15 bp, we found that 99.99%, 100%, 99.98%, and 97.95% of viral, archaea, bacteria, and eukaryotic species, respectively, had significantly fewer observed oligonucleotide kmers than expected by chance (Chi-square test with Bonferroni corrected p-value, p-value<0.05), whereas only between 0% and 0.02% cases in each of the taxonomies had higher enrichment of observed kmers per species than expected by chance. Similar results were also obtained for 16bp and 17bp kmer lengths (Figure 2b). We infer that the nullomer sequences are more frequent than expected by chance across taxonomies, and this is a universal rule across all organismal genomes likely driven by negative selection and genome repetitiveness.

### Detection of genome primes across 45,785 reference genomes

In addition to the kmer diversity and uniqueness across different taxa, we were also interested in identifying oligonucleotides that were absent from all reference genomes, which are termed as genome primes. These sequences are likely to be under selective pressure (Georgakopoulos-Soares, Yizhar-Barnea, et al. 2021; Hampikian and Andersen 2007) and have biological significance. Additionally, genome primes could serve practical purposes, such as PCR primers, highly specific platforms in CRISPR construct designs, genomic barcodes, and sample labeling. Previous studies examined a limited number of genomes and derived 60,370 fifteen bp nucleic prime sequences (Hampikian and Andersen 2007). Therefore, we aimed to determine the minimum kmer length at which we could identify genome primes with the current number of available reference genomes.

We discovered a total of 5,186,757 genome primes at sixteen bps. We characterized the genomic features of these sequences and found that they had a high GC content with average GC content of 54.33% (Figure 2c). The most prominent feature of genome primes was the presence of GC/CG dinucleotides, which occurred in 99.9997% of genome prime sequences. Only sixteen genome primes lacked GC/CG dinucleotides. Furthermore, we observed a significant enrichment of CpG sites over GpC sites. CpG sites had 1.94-fold higher frequency relative to GpC sites within genome primes (Figure 2d; **Supplementary Figure 2**; binomial test, p-value<e-100). These results suggest that genome primes are highly enriched in CpGs, possibly due to their higher mutation rate (Sved and Bird 1990; Fryxell and Moon 2005).

### Detection of nucleic quasi-primes across 45,785 reference genomes

Next, we examined if there is a certain kmer length at which we can identify sequences that are unique to a single species and absent from every other species examined, which we have defined as nucleic quasi-primes. We found that up to fifteen bps in length, every possible kmer was found in two or more species among all the genomes examined. Therefore, we concluded that nucleic quasi-primes cannot be found up to fifteen bp lengths in any genome. However, at sixteen bps we report 14,678,002 nucleic quasi-primes. We report that the number of nucleic quasi-primes detected varies substantially between species, and the median number of nucleic quasi-prime sequences detected is sixteen across the studied organisms, with certain organisms not having any nucleic quasi-prime at that kmer length (Figure 3a).

**Figure 3.**
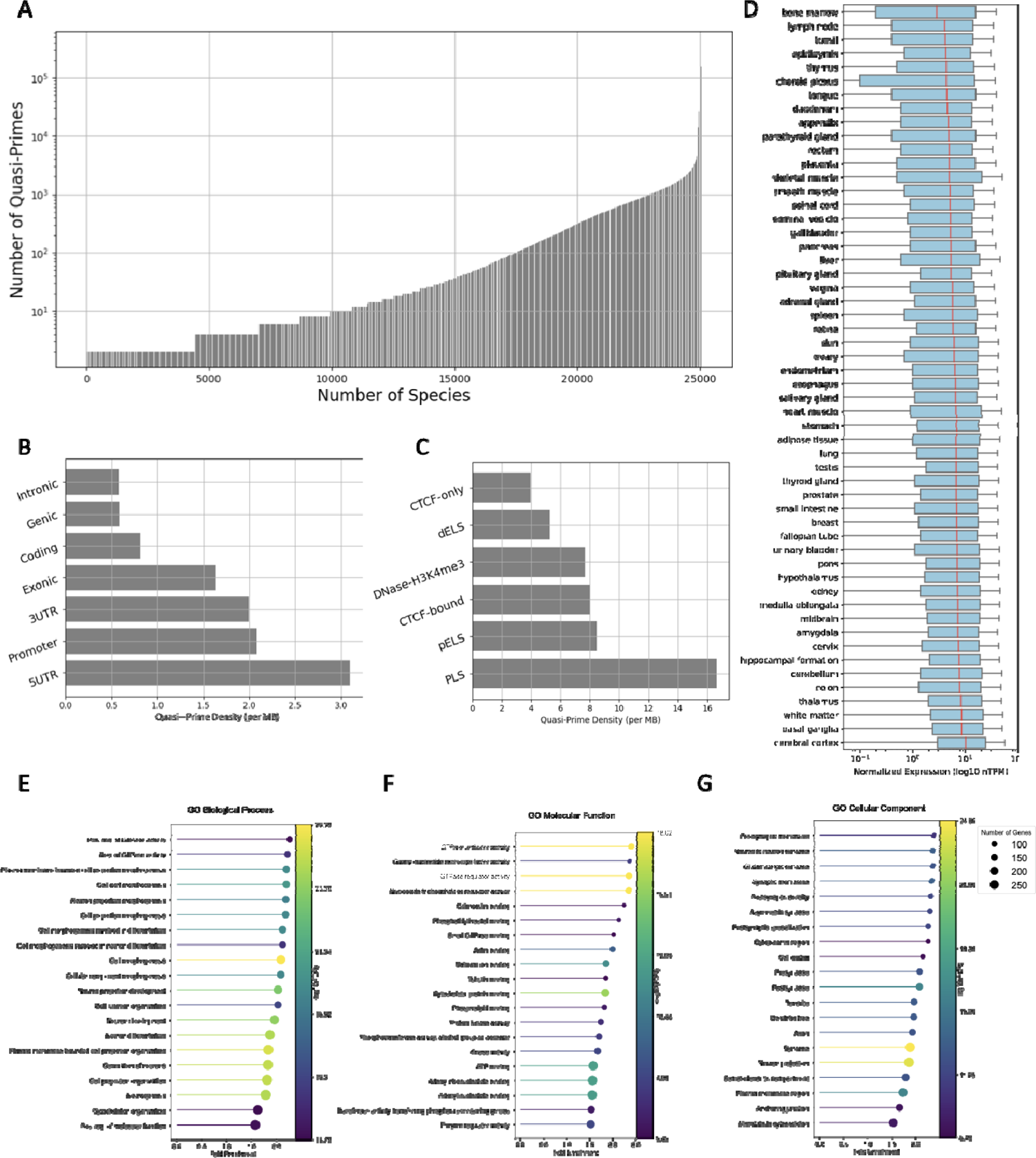
Taxonomic nucleic quasi-prime sequences. **A.** Number of quasi-primes detected in each reference genome. **B.** Density of human quasi-primes across genic regions. **C.** Density of human quasi-primes across *cis*-regulatory elements. **D.** Gene expression of human quasi-primes across tissues. **E-G.** GO term analysis for human quasi-primes for **E.** Biological processes, **F.** molecular function, and **G.** cellular components.

### Human quasi-primes are preferentially found in promoter and 5’UTR regions

Our next aim was to identify the presence and characteristics of human quasi-prime nucleotide sequences. We detected a total of 19,226 human nucleic quasi-prime sequences throughout the human genome. Analogous to genomic primes, human quasi-primes exhibited a high GC content and an enrichment of CpGs relative to GpCs (t-test, p-value<10^-100^; **Supplementary Figure 3a-c**). We also mapped the locations of each human quasi-prime sequence in the genome and identified 11,982 loci that harbor human quasi-primes of which 4,525 were in genic regions. We assessed the abundance of human quasi-primes in the following genomic sub-compartments: genic, intronic, coding, and 5′ and 3′ UTRs as well as 2,500 bp upstream of the transcription start site (TSS). We found that the highest frequency of human quasi-primes is observed at 5’UTR and promoter regions (Figure 3b). We also utilized regulatory elements as categorized by ENCODE (ENCODE Project Consortium et al. 2020) to examine the distribution of human quasi-primes across *cis*-regulatory elements. The set of *cis*-regulatory elements encompassed CTCF-only, CTCF-bound, PLS, DNase-H3K4me3, dELS, PLS, and pELS terms. We report that human quasi-prime loci are most likely to be found in PLS (Figure 3c). Thus, we infer that specific cis-regulatory elements, including promoter regions and 5’UTRs, have the highest frequency of human quasi-prime sequences.

Human accelerated regions have been previously characterized as regions that are evolutionarily conserved, but have diverged in humans (Pollard et al. 2006). Using the set of human accelerated regions collected from five previous studies (Doan et al. 2016), we examined if there is overlap with human quasi-prime loci. We find only three human quasi-prime loci overlapping human accelerated regions, accounting for 0.0156% of total human quasi-primes, indicating that human quasi-primes capture distinct genomic loci.

### Human quasi-primes are associated with brain development and cognitive function

We analyzed the expression of quasi-prime-containing genes across human tissues. In total, we examined the consensus normalized expression from RNA-seq experiments across 50 tissues (Pontén, Jirström, and Uhlen 2008). We observe that brain regions, including the cerebral cortex, basal ganglia, white matter, and thalamus show the highest expression (Figure 3d), indicating a preference for quasi-primes for brain-related genes.

Subsequently, we conducted a GO term analysis to examine the biological processes associated with human quasi-prime containing genes. We found that when examining cellular component terms, neuronal-associated terms are enriched, including synaptic and postsynaptic membrane, neuron-to-neuron synapse, dendritic and axonal terms (Figure 3g). For molecular and biological function terms, we found terms associated with GTPase activity, cell morphogenesis, and neuron morphogenesis being highly enriched (Figure 3e**,f**). Therefore, these results substantiate the brain-related roles of human quasi-prime containing genes.

### Human quasi-prime containing genes are involved in neurological diseases

We conducted an Ingenuity Pathway Analysis to assess the enrichment of quasi-prime containing genes in various biological pathways, which we will refer to as quasi-prime genes. We estimated the proportion of genes in each pathway being quasi-prime genes (ratio) and the statistical significance of the enrichment. We found that the most enriched pathways were related to “neurotransmitters and other nervous systems signaling”, such as the opioid signaling pathway (-log p-value=9.28, ratio=0.276), circadian rhythm signaling (-log p-value=7.87, ratio=0.266), dopamine-DARPP32 feedback in cAMP signaling (-log p-value=7.25, ratio=0.289), and axonal guidance signaling (-log p-value=7.245, ratio=0.224) (Figure 4a). Among the highly enriched pathways, the nervous system related pathways are found among the lowest p-values (**Supplementary Figure 4**). Other important pathways are related to cardiovascular signaling (nitric oxide signaling, cellular effects of sildenafil, cardiac hypertrophy signaling, cardiac beta-adrenergic signaling) and endocrine/neuroendocrine functions. Our results suggest that pathways associated with the nervous system are most affected by quasi-prime genes.

**Figure 4.**
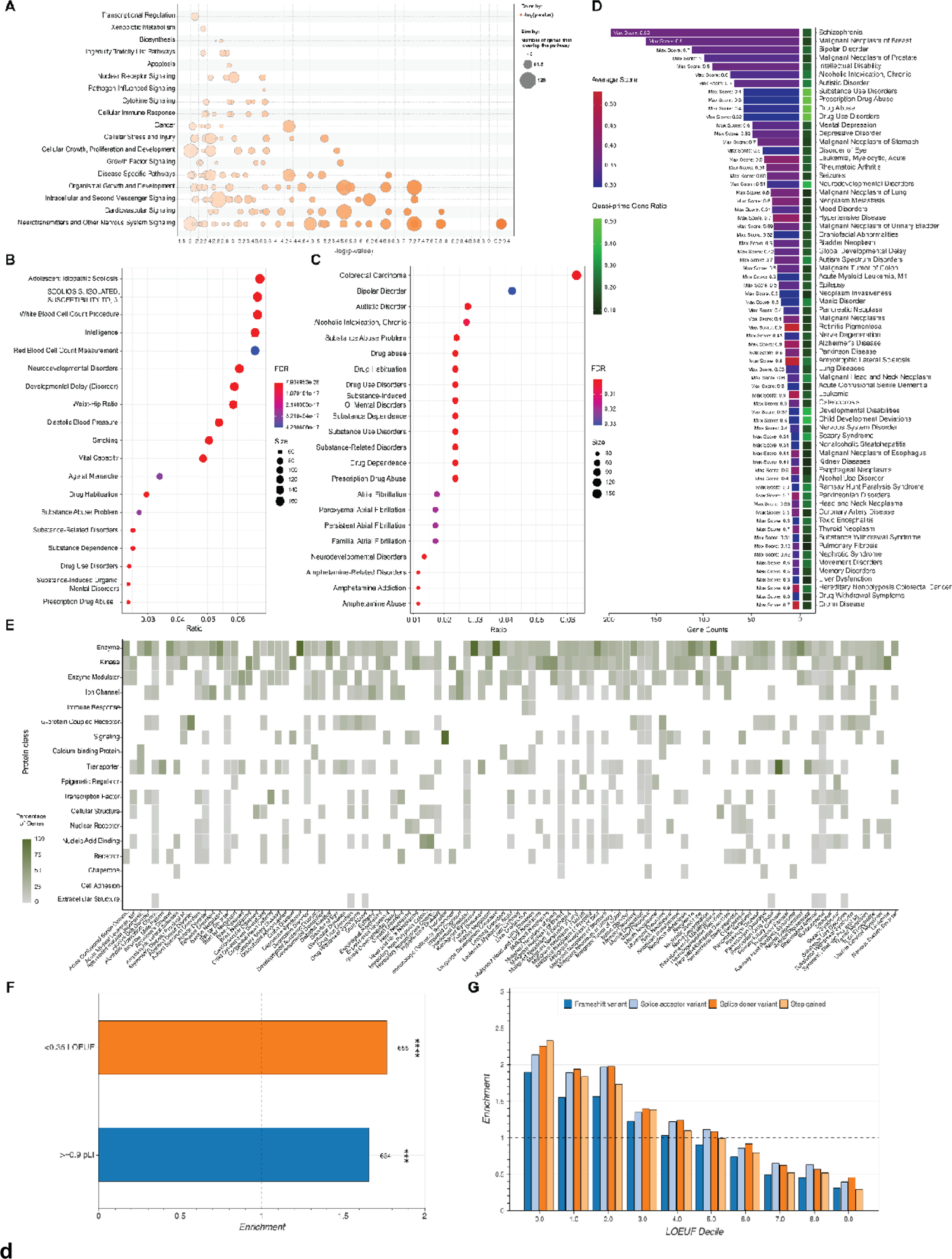
Functional and clinical disease associations of quasi-prime genes. **A.** Bubble plot showing the enriched pathway categories from IPA analysis of quasi-prime genes. The size of each bubble represents the proportion of genes in the pathway having human quasi-primes, and the color represents the adjusted p-value of the enrichment. **B.** DisGeNET enrichment dot plot showing the top characteristics associated with all quasi-prime genes present in “ALL” DisGeNET databases. **C.** DisGeNET enrichment dot plot showing the top characteristics associated with all quasi-prime genes present in the “CURATED” DisGeNET database. The dots represent the characteristic association score, and the color represents the number of genes in common between the gene set and the disease/characteristic. **D.** Gene disease bar plot presenting the count of associated genes across different disease conditions for all diseases with more than 6 associated genes. The average disease association score for all quasi-prime genes for a given disease within “ALL” DisGeNET databases is visualized in the color of the bar. The max score for all genes within a disease is displayed in text on figure. The number of quasi-prime genes out of the total genes annotated in the database is represented as a ratio shown in the heatmap tile. **E.** Protein class heat map. The color scale represents the percentage of quasi-prime genes subset within the DisGeNET “ALL” databases’ query for each disease belonging to each protein class. **F-H.** GnomAD pLOF constraint gene analysis. **F.** The odds ratio enrichment of highly constrained human quasi-prime genes (with pLI >= 0.9 and LOEUF < 0.35) is displayed as a bar plot. The enriched gene set was tested for statistical significance with a hypergeometric test. Significance levels are indicated as follows: * p < 10^-25, ** p < 10^-35, *** p < 10^-45, and **** p < 10^-55. **G.** pLOF variant type enrichment across different LOEUF Decile bins for human quasi-prime genes.

We also performed a gene-disease association analysis using DisGeNET (Piñero et al. 2017, 2020). When examining diseases and other traits that are highly enriched for human quasi-prime containing genes, we find several neurological disorders had the strongest associations, including schizophrenia, bipolar disorder, intellectual disability, drug abuse, and autism (Figure 4b**,d**). Specifically, we report that 198 schizophrenia-associated genes and 113 bipolar disorder-associated genes are also quasi-prime containing genes. We also find that the set of quasi-prime genes associated with these diseases include primarily G-protein coupled receptors, ion channels, signaling enzymes, and nucleic acid binding proteins (Figure 4e).

From an evolutionary perspective, we aimed to understand whether quasi-primes play a role of highly constrained genes and pLOF variants in disease pathogenesis. Therefore, we performed an intersection analysis of our gene set with the pLOF variant gnomAD database. Our findings revealed a significant enrichment (pLI: odds ratio: 1.66, p value: 2.65 x 10e-45, Effect Size (Cohen’s h): 0.223; LOEUF: odds ratio: 1.77, p value: 1.23 x 10e-57, Effect Size (Cohen’s h): 0.225) of highly constrained genes being quasi-prime containing genes (Figure 4f). We observed that highly constrained quasi-prime genes, as indicated by lower LOEUF deciles, were enriched (p value: 9.31a10e-08, rho:-0.987) with potentially deleterious pathogenic variants such as frameshift, splice acceptor, splice donor, or stop gained variants (Figure 4g). Our results highlight the potential role of quasi-prime containing genes in disease pathogenesis. We therefore discover disease associations with human quasi-primes, primarily those relating to cognition, with evidence to support an evolutionary role.

### Human quasi-prime genes account for a significant proportion of the variation in expression of primary motor cortex cells

We explored the contribution of quasi-prime genes to the cellular diversity of the human primary motor cortex (M1) using single-cell transcriptomics (Figure 5). We leveraged the data from the Allen Institute for Brain Science, which obtained single-cell profiles from postmortem and neurosurgical donor brains with dissected cortical layers (Bakken et al. 2021) **(Supplementary Figure 6**). We identified 442 quasi-prime genes (22.1%, p-value: 1.613×10^-173^, Effect size: 0.165, Odds ratio: 2.7) among the top 2,000 genes with the highest average expression variability across M1 cell types (Figure 5a**; Supplementary Figure 7**). The most variable quasi-prime genes were *RELN*, *THSD7B*, *FBXL7*, *GPC5*, and *ADARB2*. The most variable quasi-prime gene, *RELN,* encodes for reelin, a protein that regulates neuron signaling during brain development. Our results demonstrate that quasi-prime genes account for a significant proportion of the single-cell expression variation among M1 primary motor cortex cells.

**Figure 5:**
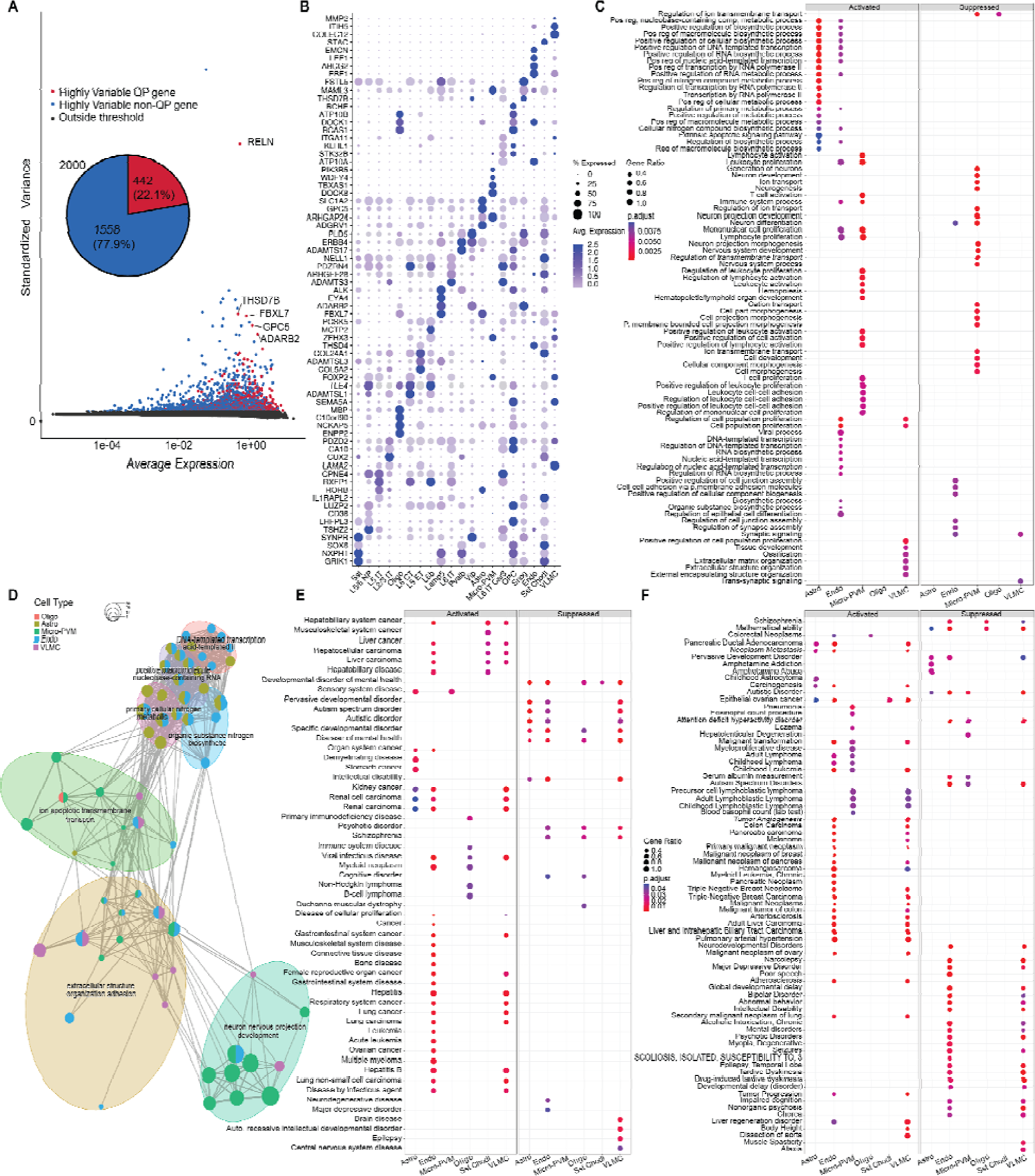
Differentially expressed genes associated with quasi-prime sequences among cell types found in the human single-cell primary motor cortex brain atlas. **A.** 2,000 genes were thresholded by variation to capture the highly variable genes. The resulting highly variate genes were labeled either as genes with quasi-prime loci or without. 442 genes associated with quasi-prime loci (22.1%) capture high levels of variation as shown in the pie chart (Fisher’s exact test p-value: 1.613 x 10^-173^, Effect size: 0.165, Odds Ratio: 2.7). **B.** Differential expression of quasi-prime genes thresholded by an absolute log2 fold change > 1 and p adjusted value < 0.05 represented by a dot plot. The top four most significant genes per above cell type ranked by positive log2 fold change are shown. **C.** Differentially expressed quasi-prime genes were utilized in gene set enrichment analysis of the gene ontology database representing the top twenty terms per category thresholded by a p-value < 0.01. **D.** A cmap network graph shows the clustering of top categories in the GO term GSEA analysis. **E-F.** Significant (p value < 0.05) differentially expressed genes with an absolute log2 fold change > 1 amongst cell types were used in GSEA analysis as represented by a dot plot. **E.** Disease ontology database. **F.** DisGeNET “ALL” database.

### Human quasi-prime genes are linked to neuronal support and protection in the primary motor cortex among non-neuronal cells

Differential expression analysis across cell types was conducted to determine cell-type specific expression patterns and function using quasi-prime genes. Our analysis revealed non-neuronal cell types including astrocytes, endothelial cells, microglia perivascular-macrophages, oligodendrocytes, and vascular and leptomeningeal cells exhibited the most distinct expression profiles of quasi-prime genes with cell-type specific markers (Figure 5b**; Supplementary Figure 8**). We performed a gene ontology (GO) gene set enrichment analysis (GSEA) of the most significantly differentially expressed quasi-prime containing genes (log2 fold change > 1 and p-value < 0.05) for each of the twenty cell types. We observed that the quasi-prime gene sets of the five non-neuronal cell types (astrocytes, endothelial cells, microglia perivascular macrophages, oligodendrocytes, and vascular leptomeningeal cells) had common characteristics. These cell types showed upregulation of quasi-prime gene sets involved in metabolic processes, cellular senescence, cell adhesion and proliferation, and immune system pathways (Figure 5c**,d**). They also showed downregulation of quasi-prime gene sets related to cell/tissue development, organization, signaling, and transport. Among these cell types, microglia perivascular macrophages exhibited enhanced expression of quasi-prime gene sets associated with immune development, proliferation, adhesion, and activation and reduced expression of quasi-prime gene sets associated with neuro- and morpho-genesis, and neuronal and cell development. Oligodendrocytes exhibited reduced expression of quasi-prime gene sets involved in ion transmembrane transport regulation pathways. These results suggest that the human quasi-prime gene sets in the primary motor cortex are predominantly expressed in non-neuronal cell types that regulate the cell and tissue environment to support and protect neuronal cell development.

### Human quasi-prime genes in non-neuronal cells of the primary motor cortex are associated with neurological, behavioral diseases and cancer

Furthermore, we performed GSEA of differentially expressed quasi-prime genes for cell-type specific disease association analysis using two additional databases: disease ontology (DO) (Figure 5e) and DisGeNET (Figure 5f). The GSEA results from both databases were consistent and confirmed our previous findings of cognitive, developmental, behavioral, and cancer associations. We found that the activated biological pathways associated with quasi-prime genes were mainly associated with cancer, immune system diseases, and viral infectious diseases in the five non-neuronal cell types. These cell types also had underexpressed biological pathways associated with substance abuse and addiction and cognitive, mental, and developmental disability and disorder. Astrocytes had overexpressed biological pathways involved in sensory system and demyelinating diseases as well as underexpressed biological pathways associated with substance addiction and abuse. Microglia perivascular macrophages had activated biological pathways associated with immune related cancers and viral infectious disease. Interestingly, these cell types had underexpressed biological pathways linked to attention deficit disorder, autism, and schizophrenia while oligodendrocytes also had suppressed biological pathways linked to schizophrenia and mathematical ability. We also found associations of quasi-prime genes in two neuronal cell types: somatostatin chondrolectin (Sst Chondl), a GABAergic interneuron; and layer 5 extra telencephalic-projecting neurons (L5 ET), a type of glutamatergic neuron found in the neocortex.

Two clusters of cognitive/mental/developmental diseases and cancer/carcinomas among the network graphs of each disease database are visualized **(Supplemental Figure 9**). These diseases are known to be associated with the non-neuronal biological pathways of neuronal development, support, and protection derived from gene ontology GSEA. Therefore, human quasi-prime gene sets are most associated with two disease classes and are useful to determine cell specific mechanistic relationships of biological significance for cognitive/mental/developmental disease and cancer.

### Human nucleic quasi-primes are crucial determinants of disease variant distribution and quantitative trait loci

An analysis of gnomAD v2 variants revealed that human quasi-primes exhibit a greater likelihood to be a constrained gene than would be expected by random chance. This is substantiated by thresholds of elevated constraint (pLI >= 0.9: p-value: 2.65×10−45, odds ratio: 1.66, effect size (Cohen’s h): 0.22; and LOEUF 90% upper < 0.35: p-value: 1.23×10−57, odds ratio: 1.77, effect size (Cohen’s h): 0.26) (Figure 5f). Furthermore, we observed an enrichment of potential loss-of-function (pLOF) variant types such as frameshift (Decile bin: 0-4), splice acceptor (Decile bin: 0-5), splice donor (Decile bin: 0-5), and stop gained (Decile bin: 0-4) variants for human quasi-primes across a diverse range of constraint as indicated by the LOEUF Decile (Figure 5h). Our data strongly suggest that human quasi-primes are overrepresented in highly constrained regions with an increased number of pathogenic and pLOF variants (Spearman’s correlation: rho: −0.99, p-value: 9.31×10^−8^), further providing evidence towards their potential role in genetic diseases.

We investigated the likelihood of variants identified through Genome-Wide Association Studies (GWAS). The risk of disease or specific traits exhibit a greater likelihood to coincide with or to be located in regions near human quasi-prime loci. We generated a simulated control human quasi-prime set containing sequences within 10kB from the original locus and had the same GC content for comparison. Our findings indicate that variants derived from GWAS exhibit a 3.78-fold increased likelihood of directly overlapping human quasi-prime loci compared to matched controls (Binomial test, p-value<2*10^-13^) and show a 4.43-fold enrichment relative to the background vicinity (Figure 6a). These results suggest that human quasi-prime sequences possess an increased presence of variants that exhibit substantial associations with human diseases and traits.

**Figure 6:**
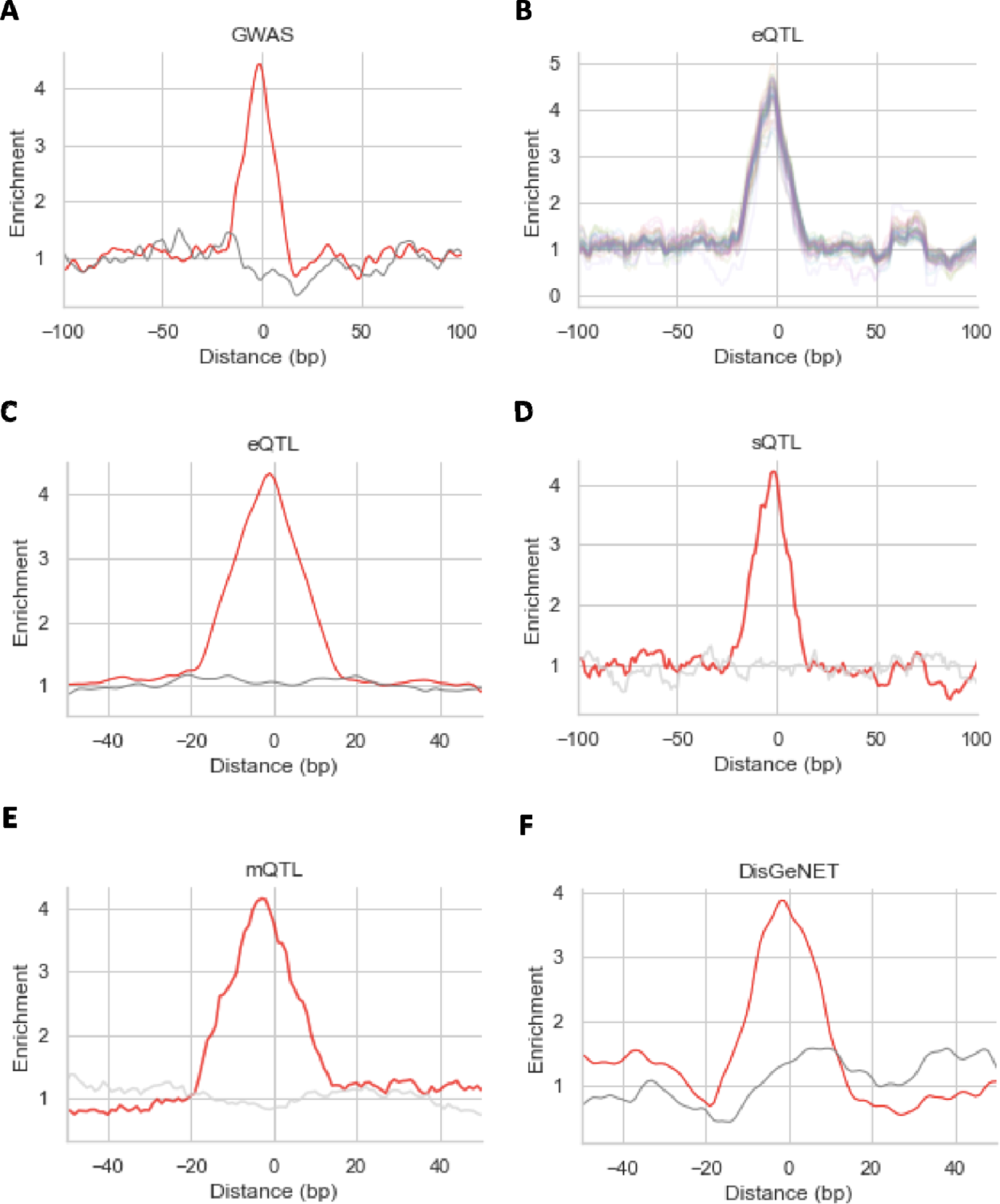
Analysis of disruptive and regulatory variants. **A.** Enrichment of human quasi-primes at and around GWAS variant loci. **B.** Enrichment of human quasi-primes at and around eQTL loci. Each line represents a tissue. **C.** Enrichment of human quasi-primes at and around multi-tissue eQTL loci. Enrichment of human quasi-primes at eQTL loci. **D.** Enrichment of human quasi-primes at and around sQTL loci. **E.** Enrichment of human quasi-primes at and around mQTL loci. **F.** Enrichment of human quasi-primes at and around DisGeNET-derived loci. Gray lines represent the enrichment for simulated controls of human quasi-prime.

Additionally, we examined expression quantitative trait loci (eQTL) from 49 tissues using GTEx (GTEx Consortium 2020) to provide additional evidence for the functional roles of human quasi-prime loci. We observe that across the tissues examined, single tissue eQTLs are enriched at human quasi-prime loci relative to controls (t-test, p-value=3.5*10^-85^). Across the tissues examined, there is on average a 4.34-fold enrichment of eQTLs relative to the surrounding regions (Figure 6b), and they are 3.92 times more likely to overlap the quasi-prime loci relative to the matched controls (Binomial test, p-value<5*10^-8^). Similar results were observed when examining eQTLs found across multiple tissues, reporting a 4.35-fold enrichment at human quasi-prime sequences over surrounding regions (Figure 6c).

We next examined if human quasi-primes are also enriched for genetic variants that affect alternative splicing or methylation. To that end, we analyzed splicing QTLs (sQTLs) across the 49 tissues available in GTEx. We find that across the examined tissues, sQTLs are 4.75-fold more likely to overlap human quasi-prime loci than their matched controls (t-test, p-value=4.67*10^-13^) and are 4.29-fold enriched over their surrounding sequences (Figure 6d). Methylation QTLs were also examined across nine tissues and we observed 5.37-fold more mQTLs at human quasi-primes relative to matched controls (t-test, p-value<0.001) (Figure 6e). We therefore conclude that expression, splicing and methylation QTLs are significantly enriched at human quasi-prime loci.

DisGeNet provides curated variant-disease associations derived from multiple sources (Piñero et al. 2017, 2020). We examined whether variants associated with human diseases are enriched at human quasi-prime sequences. Human quasi-prime loci are observed to be 2.5-fold more likely to overlap with disease variants than expected by chance using the simulated control loci (Fisher two-sided test, p-value< 0.001), and they are 3.86-fold enriched over surrounding regions (Figure 6f). These results further validate our previous associations and provide support for the importance of studying human quasi-primes to gain insights into human traits and diseases.

## Discussion

In this work, we have analyzed 45,785 reference genomes and identified the shortest nucleic kmers that are unique to each species’ genome, which we refer to as nucleic quasi-primes. This extends our recent work on the detection of peptide quasi-primes (Mouratidis et al. 2023) and also provides multiple biological insights and potential future applications. We also identify primes, the set of shortest kmers that are absent from all studied genomes, which, similarly to quasi-primes, exhibit an unusually high GC content and are enriched for CpGs. Because CpGs are hypermutable (Sved and Bird 1990; Fryxell and Moon 2005), we postulate that their absence is at least in part driven by the higher mutation rate. Further examination of genome primes in future studies and investigation of potential biological functions would be of high interest, especially to understand if their presence is detrimental to organismal fitness.

We perform a case study on the set of human nucleic quasi-primes and find that they are enriched in genes that are highly expressed in regions associated with brain development and function. We also discover associations between quasi-prime containing genes and neurological disease associated genes, including schizophrenia, bipolar disorder, intellectual disability, autism, addiction, and drug abuse. These findings are supported by the expansion of brain size and cognitive tasks following the divergence of humans from other primates (Florio, Borrell, and Huttner 2017). The faster turnover rate of CpG sequences within human quasi-primes can result in evolutionary adaptations and can account for human-specific traits and molecular phenotypes. Characteristics of schizophrenia, which is the disease with the strongest association with human quasi-primes, have not been observed in other species (Burns 2004). Similarly, a strong association of scoliosis, a bipedal specific disorder (de Reuver et al. 2021), and hepatitis/hepatitis B, viral disease derived from a humanoid host pathogen (Devaux et al. 2019), were observed (Figure 4) indicating that human quasi-primes can identify species-specific traits. Identification of quasi-primes can therefore provide insights into the acquisition of new traits and identify regions of accelerated evolution.

Similarly to nucleic primes, human quasi-primes have high GC content and CpG sites. Additionally, human quasi-primes are enriched for mQTLs. We therefore speculate that the presence of epigenetic changes found within human nucleic quasi-primes could partially account for their human-specific roles. Genes within the human genome exhibiting a high degree of constraint are predisposed to being classified as quasi-prime genes. Furthermore, highly constrained quasi-prime genes demonstrate a significant enrichment for variants that are pathogenic and potentially lead to loss of function. In future studies, it will be of interest to use nucleic quasi-primes to further explore and understand the functional genomic elements that are linked to phenotypic changes and the evolution of species-specific traits.

Quasi-prime loci can also be utilized to gain insights in cell-type specific associations. In a human case study of the primary motor cortex at single cell resolution, we highlight biological and disease associations among brain cell types to functionally profile individual cell types via human quasi-prime genes. Genes containing quasi-primes account for noteable gene expression diversity and heterogeneity across cells in the primary motor cortex. This heterogeneity may be crucial for the tissue’s overall function, enabling it to respond to diverse stimuli or perform specialized tasks. Interestingly, the most variable quasi-prime genes are related to neuronal signaling, tissue reorganization, transferase activity, proteoglycan binding, and RNA-editing activity. It’s worth noting the activation of biological pathways related to cellular senescence, cell proliferation, and immune system pathways and the suppression of pathways related to cell/tissue development, organization, signaling, and transport among the most variable expressed genes in non-neuronal cells. We therefore indicate that unique species-specific gene sets can deliver cell-specific information highlighting a connection to heritable disease mechanism. We also observe that human quasi-primes are hotspots for pathogenic variants and for variants that significantly alter gene expression, or result in alternative splicing or variable DNA methylation. This can lead to further research regarding evolutionary adaptations in humans and can enable an improved understanding of human diseases.

Additionally, several applications using nucleic quasi-primes are possible and can be the target of future studies. One of these is the usage of nucleic quasi-primes for the high-throughput detection of multiple organisms in diverse settings, including pathogen detection in clinical settings or in food safety applications such as foodborne outbreaks from microbial contamination (Tringe and Rubin 2005). Nucleic quasi-primes could also enable the detection of invasive or rare species with consequences in conservation. Detection based on nucleic quasi-primes could be coupled with other existing technologies including CRISPR nucleases that cut in species-specific genomic loci (Kellner et al. 2019) or adaptive sequencing, which enriches for a set of short sequences (Loose, Malla, and Stout 2016). Additionally, population variants that disrupt or introduce human quasi-primes could be used as a unique identification signature of individuals and could result in a new type of DNA fingerprinting method. Finally, genome primes are also of high practical interest. Applications can include their usage as genetic barcodes to track cells or organisms. Another potential application may involve their usage as highly-specific CRISPR-Cas9 landing pads in genetic engineering and therapeutic applications.

Finally, as the representation of genomic diversity in nature increases with the sequencing of more organismal genomes, the identification of nucleic quasi-primes will provide succinct genomic fingerprints for every available organism.

## Materials and Methods

### Reference genomes used

Collection of reference genomes was performed for the GenBank and RefSeq databases as well as 104 reference genomes from the UCSC genome browser website (Supplementary Table 2). Reference genomes present in multiple databases were deduplicated. For each genome in each database kmer extraction was performed for kmer lengths up to 16bp, as previously described in (Georgakopoulos-Soares, Yizhar-Barnea, et al. 2021). Only DNA letters “A”, “C”, “G”, and “T” were considered across the analyses, and other IUPAC letters including “N” were ignored. Quasi-prime nucleotides were defined as nucleotide sequences found in only one reference genome, which were absent in every other genome analyzed.

### Genome simulations

Simulated genomes were generated using Ushuffle (Jiang et al. 2008) for every reference genome controlling for dinucleotide content. For each simulated genome oligonucleotide kmer extraction was performed for kmer lengths up to and including 17bp and compared against the number of kmers identified in the reference genomes (Figure 2b). For each reference genome and each simulated genome, the number of kmers identified was compared, and significance was estimated using binomial tests with Bonferroni corrections.

### Nucleic quasi-prime definition

Let us define the DNA alphabet *L* = {*A*, *T*, *C*, *G*}. We define a DNA sequence of length n as A=a1 a2a3…an□, where ai∈L. We denote its reverse complement sequence A’=an’a_n-1’’…a2’a1’□, where ai’= A if ai=T, ai’=G if ai’=C, ai’ =Cif ai=G and ai’=T if ai=A. A genome GN is a set of DNA sequences GN = {Ai | i =1,…, m} U { Ai’ | i = 1,…,m}. We define C = {GN1, GN2, GN3, …, GNk} the set of the k genomes analyzed, where GNi is the genome of the i-th species analyzed.

For this paper, a DNA kmer is defined as a short DNA sequence K=k1k2…kl with length l ∈ N ^ l <= 16. We define K ∈ G if and only if there is DNA sequence A = a_1a_2a_3…a_n□ ∈ G and 1 <= i <= n-l, such as a_ia_i+1…a_i+l-1 = k1k2. kl. We developed an algorithm in Python that performs an exhaustive search across input sequences A, identifying all kmers such as K ∈ G. We define the set of quasi-primes of a genome Gi as Q = {K | K ∈ Gi ^ q ∉ Gj for j!=i}.

### Quasi-prime genomic analyses

Identification of the genomic locations of quasi-primes was performed using a custom Python script. Gencode annotation (v43) was used to derive gene information (Frankish et al. 2023). Genomic sub-compartments namely, genic regions, intronic regions, coding regions, and 5′ and 3′ UTRs as well as 2,500 bp regions upstream of the transcription start site (TSS) were derived with the UCSC Table Browser (Nassar et al. 2023). *Cis*-regulatory elements as defined by ENCODE were used for the analyses (ENCODE Project Consortium et al. 2020). *Cis*-regulatory elements included CTCF-only, CTCF-bound, PLS, DNase-H3K4me3, dELS, and pELS terms. BEDTools utilities v2.21.0 (Quinlan 2014) were used to perform the analyses and estimate the density of human nucleic quasi-primes across each genomic element (Figure 3b-c). Human accelerated regions were derived from (Doan et al. 2016) and liftOver was used to transfer them to hg38 reference genome coordinates. The intersection between human quasi prime genomic loci and human accelerated regions was performed with the function intersect from BEDTools (Quinlan 2014).

### Bulk RNA-seq analysis

Consensus normalized expressions were downloaded from The Human Protein Atlas (Pontén, Jirström, and Uhlen 2008) (RNA consensus tissue gene data) to plot expression levels of quasi-prime genes across 50 tissues. The downloaded data was based on The Human Protein Atlas version 23.0 and Ensembl version 109 (Martin et al. 2023) (Figure 3d).

### GO term analysis

A GO term analysis was performed for genes that contained at least one human quasi-prime sequence in the reference human genome with ShinyGO for GO Biological Process, GO Molecular Process and GO Molecular Function (Figure 3e-g) (Ge, Jung, and Yao 2020).

### Ingenuity Pathways Analysis (IPA)

The list of quasi-prime genes was uploaded to the IPA platform (Qiagen, Redwood City, CA) and a pathway analysis for human species only was performed (Figure 4a). The significance values (p-value of overlap) for the pathways was calculated by the right-tailed Fisher’s Exact test.

### DisGeNET enrichment analysis

The 2,492 quasi-prime genes were used in a DisGeNET enrichment analysis (Piñero et al. 2020) to determine which diseases the list of genes is most associated with the “CURATED” (Figure 4b) and the “ALL” (Figure 4c) databases using the disease_enrichment(vocabulary=”HGNC”) method. The Unified Medical Language System (UMLS) (Bodenreider 2004) was used to identify relevant diseases to query in the disease2gene() method from DisGeNET. The “MRCONSO.RRF” file was downloaded from the UMLS database. The file was filtered to retain values from the english language and MSH database while removing duplicated CUI values and any disease names with numbers. This database was subset to reflect disease names considering relevant keywords **(Supplemental Table 1)**. Additional CUIs were added to reflect diseases, disease classes, and potential human specific diseases of interest **(Supplemental Table 1)**. The resulting list of CUIs were split into multiple disease2gene() DisGeNET queries as the max allowed per query at this present time is 447 diseases. A cutoff of 0.3 was used when querying the DisGeNET “ALL” database. The resulting DisGeNET S4 classes were combined and were subsequently stratified to reflect disease enrichment results for genes associated with quasi-prime regions. The count of individual quasi-prime associated genes per disease were calculated and displayed as a bar plot (Figure 4d). A ratio of the amount of quasi-prime associated genes per disease and the total genes associated with a disease was calculated. Diseases with six or less associated quasi-prime genes were filtered from this analysis. A heatmap representing the class of proteins that the genes located in quasi-prime regions are most associated with per disease was subsequently generated.

### Single-cell RNA-seq analyses

The human M1 primary motor cortex dataset, available from Allen Institute for Brain Science (Bakken et al. 2021), was used to analyze the relationship of quasi-prime defined genes from GO term analysis among cell types found in the brain. The human M1 primary motor cortex dataset was analyzed using the seurat (version 4.3.0.1) package in R (Hao et al. 2021; Stuart et al. 2019; Butler et al. 2018; Satija et al. 2015) **(Supplementary Figure 6**). The data was normalized using the shifted logarithm (Comparison of normalization strategies can be found in **(Supplementary Figure 6a**), FindVariableFeatures() was used to generate the 2000 most variable genes (Figure 5a), and the data was scaled using scaleData(). FindAllMarkers(features=”quasi-prime-genes”, thresh.use=1, min.pct=0.25) was used to calculate the differentially expressed quasi-prime genes (DEGs) between each cell type. DEGs were thresholded by p-adjusted value < 0.05 and used for analysis (Figure 5b; **Supplementary Figure 7**). The resulting DEGs were converted to Entrez format and used in gene set enrichment analysis of Gene Ontology (Figure 5c,d), DisGeNET, and Disease Ontology (Figure 5e**,f**) databases using the comparecluster() method in the ClusterProfiler package (version 3.17) (Wu et al. 2021; Yu et al. 2012). Gene sets with a minimum of 3 genes were included in each analysis.

### Variant analysis

The GWAS catalog was derived from https://www.ebi.ac.uk/gwas/api/search/downloads/full (MacArthur et al. 2017). Multi-tissue and single tissue eQTLs were derived from the GTEx consortium (v8) (GTEx Consortium 2020). sQTLs were derived from the GTEx consortium (Garrido-Martín et al. 2021; GTEx Consortium 2020). mQTLs were derived for nine tissues from the GTEx consortium (GTEx Consortium 2020; Oliva et al. 2023). For the mQTL analysis only events with q-value<0.05 were analyzed. DisGeNET variants were derived from https://www.disgenet.org/static/disgenet_ap1/files/downloads/variant_associations.tsv.gz (Piñero et al. 2020). A set of simulated control human quasi-primes were generated; in each human quasi prime a locus that was within 10kB from the original locus and had the length and GC content was randomly selected. Enrichment for genomic variants was calculated as described in (Georgakopoulos-Soares et al. 2018) (Figure 6).

We conducted a variant analysis to investigate the enrichment of quasi-prime regions within variant regions in RegulomeDB (Dong et al. 2023). We simulated control regions to compare the potential enrichment. The Bioframe python package (v0.4.1) (Open2C et al. 2022) was used to determine the overlap between quasi-prime regions and RegulomeDB regions and control regions and RegulomeDB regions, respectively. A binomial test was then used to test for enrichment between the groups (SciPy v1.11.2). The enrichment (represented as odds-ratio) of each variant characteristic for these overlapped variant groups from RegulomeDB were subsequently plotted **(Supplementary Figure 5a,b**).

The Genome Aggregation Database (gnomAD) constraint database (Chg38) was downloaded from: https://storage.googleapis.com/gcp-public-data--gnomad/release/2.1.1/constraint/gnomad.v2.1.1.lof_metrics.by_gene.txt.bgz (Karczewski et al. 2020) and subsequently queried to evaluate the enrichment (expressed as odds-ratio) of human quasi-prime genes within regions of high constraint. A hypergeometric enrichment test was conducted to compute a p-value and effect size for genes deemed highly constrained (pLI => 0.9 or an oe_lof_upper < 0.35) (Figure 4f).

All predicted loss-of-function (pLOF) variants were obtained from gnomAD (Karczewski et al. 2020): https://storage.googleapis.com/gcp-public-data--gnomad/papers/2019-flagship-lof/v1.0/gnomad.v2.1.1.all_lofs.txt.bgz. The data frames were merged based on the position of the variant within the constraint/genic region. The enrichment of quasi-prime genes, based on LOEUF Decile (Karczewski et al. 2020) bins for each variant type, was also graphically depicted (Figure 4g). A Spearman correlation analysis was conducted to examine any potential correlational relationship between the constraint metric LOEUF Decile and variant types as a sum for quasi-prime enrichment.

## Data Availability

The quasi-prime sets derived are available at www.kmerdb.com, where we have developed a browser for the dataset. We provide a downloadable analysis of detailed frequency and annotation information for each species.

## Code Availability

All code to perform case study analysis is provided at https://github.com/Georgakopoulos-Soares-lab/quasi-prime-humancasestudy.

## Contributions

IM and IGS conceived the concept of DNA quasi-primes. IGS supervised the work. IM, MAK, NC, CC, DVC, and IGS wrote the code. IM, MAK, NC, CC, DVC, and IGS performed the analyses. MAK, CC, DVC and IGS generated the visualizations. IM, MAK, NC, CC, and IGS wrote the manuscript with help from all other authors.

## Acknowledgments

This study was funded by the startup funds of IGS from the Penn State College of Medicine and by the Huck Innovative and Transformational Seed Fund (HITS) award from the Huck Institutes of the Life Sciences at Penn State University.

## Supplementary Material

**Supplementary Figure 1:**
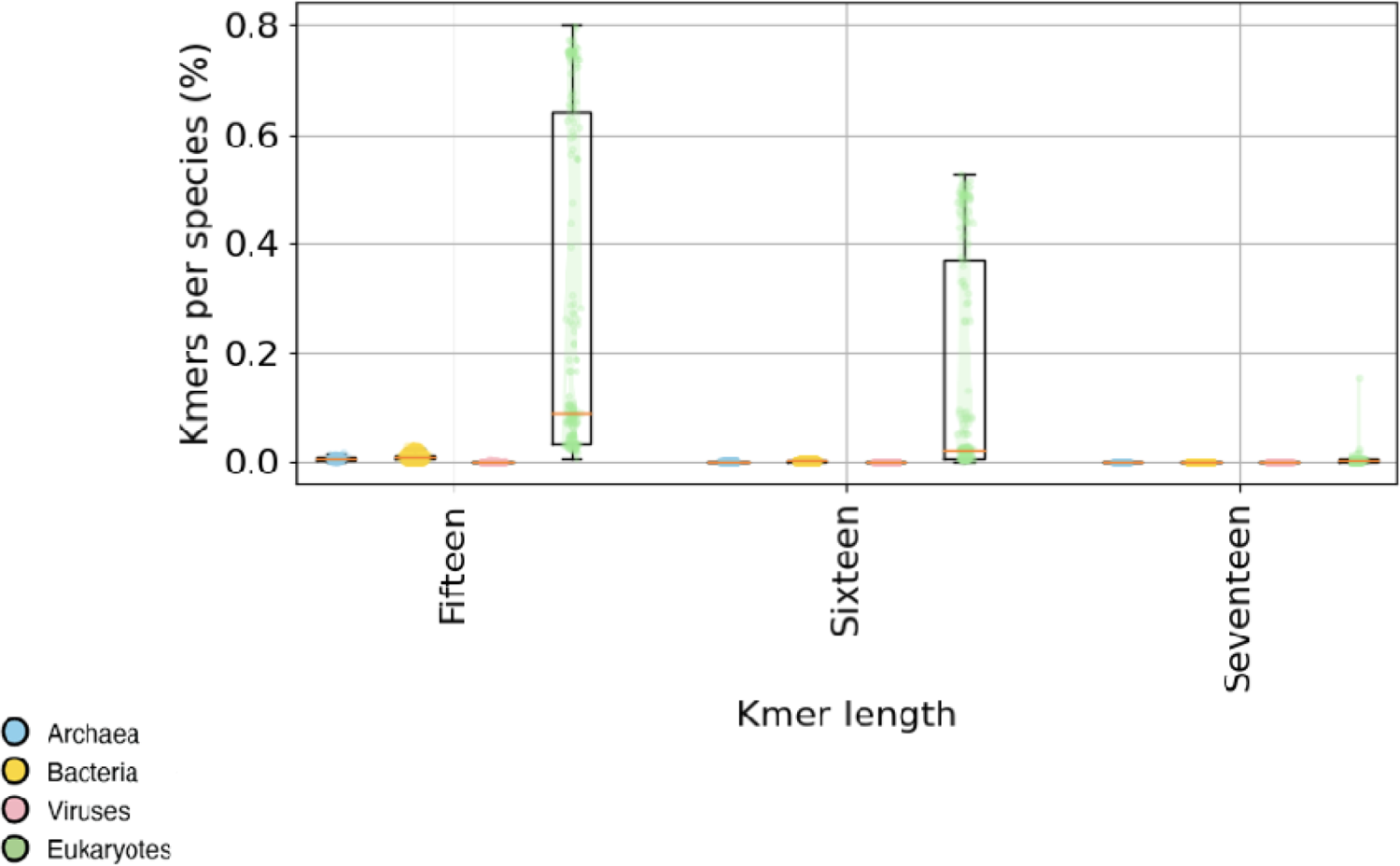
Proportion of kmers found in each reference genome for kme lengths of 15bp, 16bp, and 17bp across the taxonomic subdivisions. Each dot represents the proportion of kmers observed in a reference genome. The majority of kmers are found in a minority of the species studied across taxonomies. Error bars represent standard deviation.

**Supplementary Figure 2:**
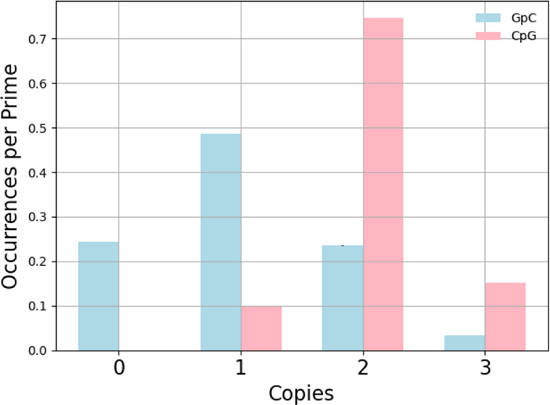
Number of occurrences of GpCs and CpGs for different copy numbers, per nucleic prime sequence. Error bars are derived from bootstrapping with replacement (n=1,000) and represent standard deviation.

**Supplementary Figure 3:**
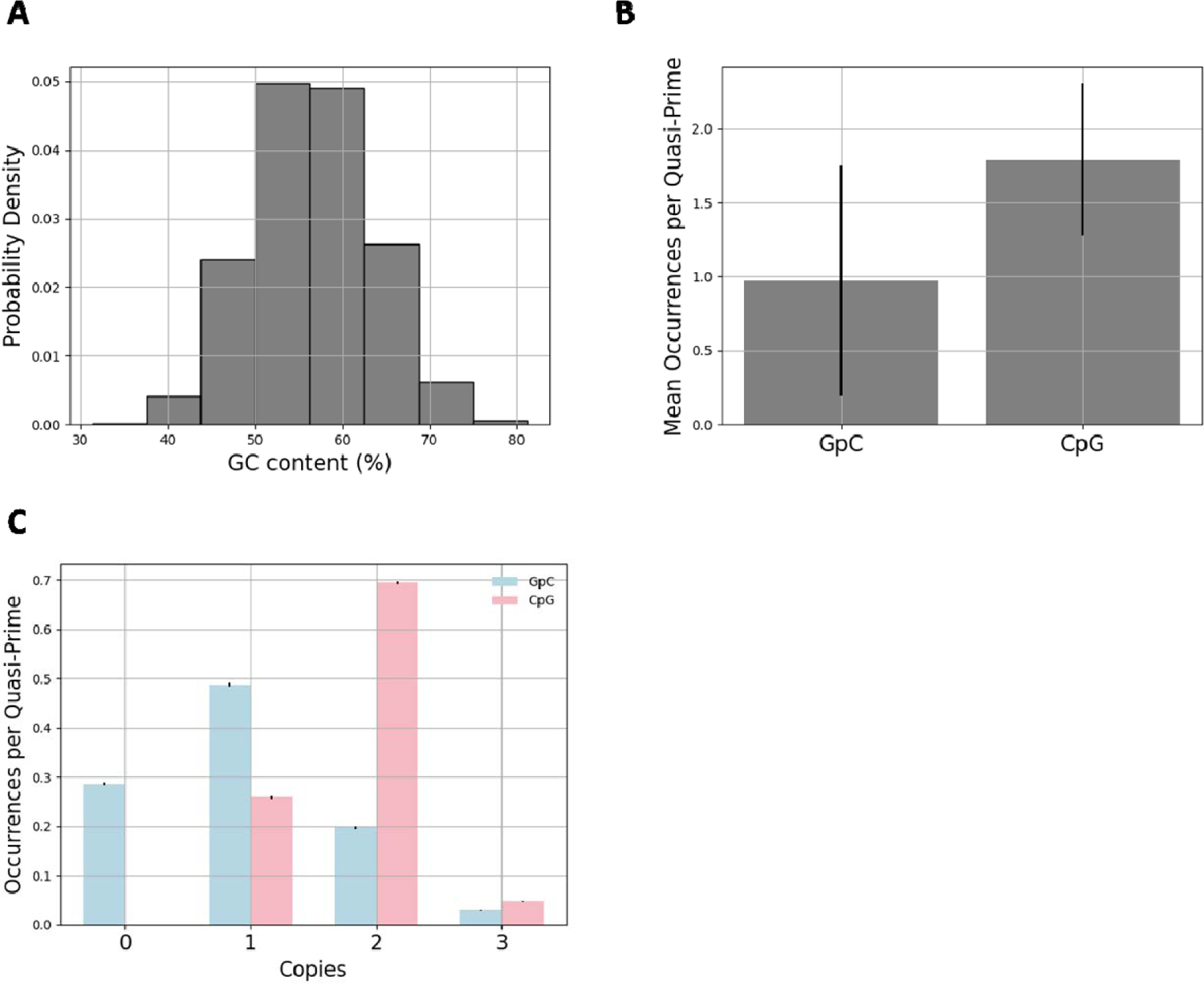
GC content distribution of human quasi-prime sequences. **A.** GC content percentage of human quasi-prime sequences. **B.** Average number of GpC and CpG occurrences per human quasi-prime. Error bars show standard deviation. **C.** Number of occurrences of GpCs and CpGs for different copy numbers, per nucleic quasi-prime sequence. Error bars are derived from bootstrapping with replacement (n=1,000) and represent standard deviation.

**Supplementary Figure 4:**
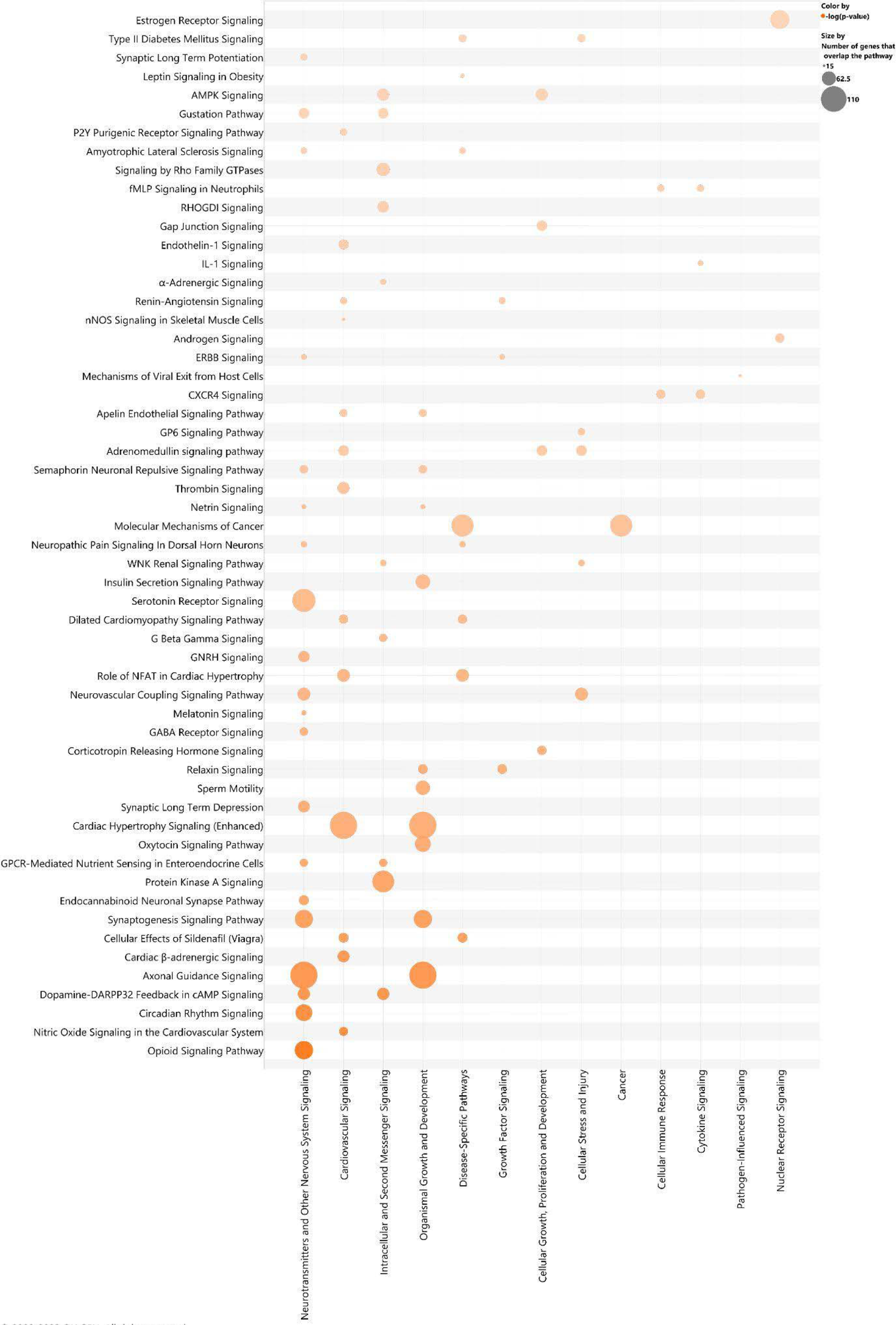
Bubble plot showing the pathway categories (Y axis) of the enriched canonical pathways (X axis) enriched in the quasi-prime gene set. The color represents the adjusted p-value of the enrichment (Increasing orange hue, the lower the p-value. The cutoff of p-value was set at <0.001). The size of each bubble represents the number of overlapping genes in the pathway.

**Supplementary Figure 5:**
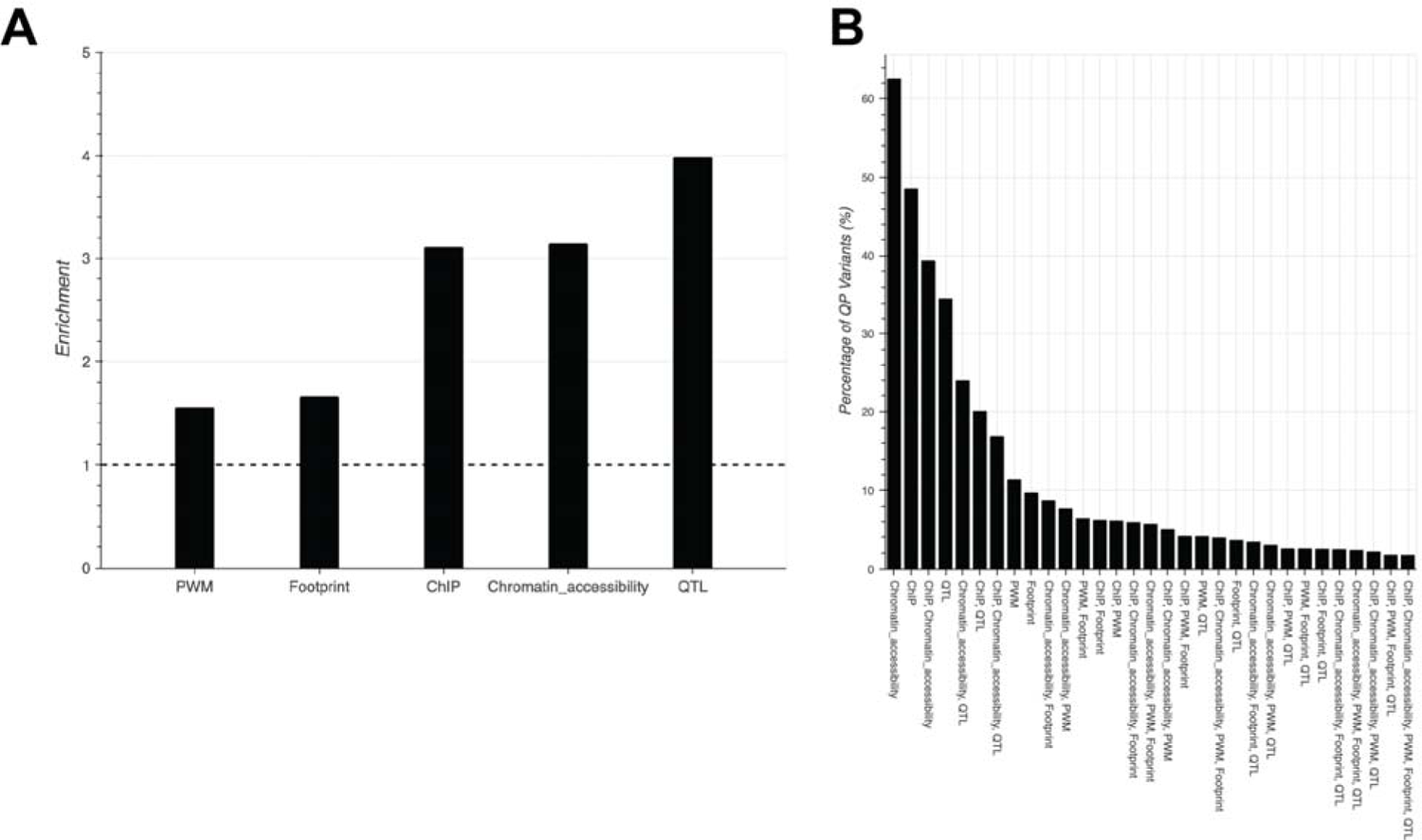
Variant Characterization of Human Quasi-primes. **A.** Enrichment (odds ratio) of quasi-prime regions as compared to simulated controls within the regulomeDB database displayed as bar plot (Binomial test p=0.0) **B.** The percentage of quasi-prime variants for different combinations of how a variant is characterized or where it is located from regulomeDB is displayed as a bar plot.

**Supplementary Figure 6:**
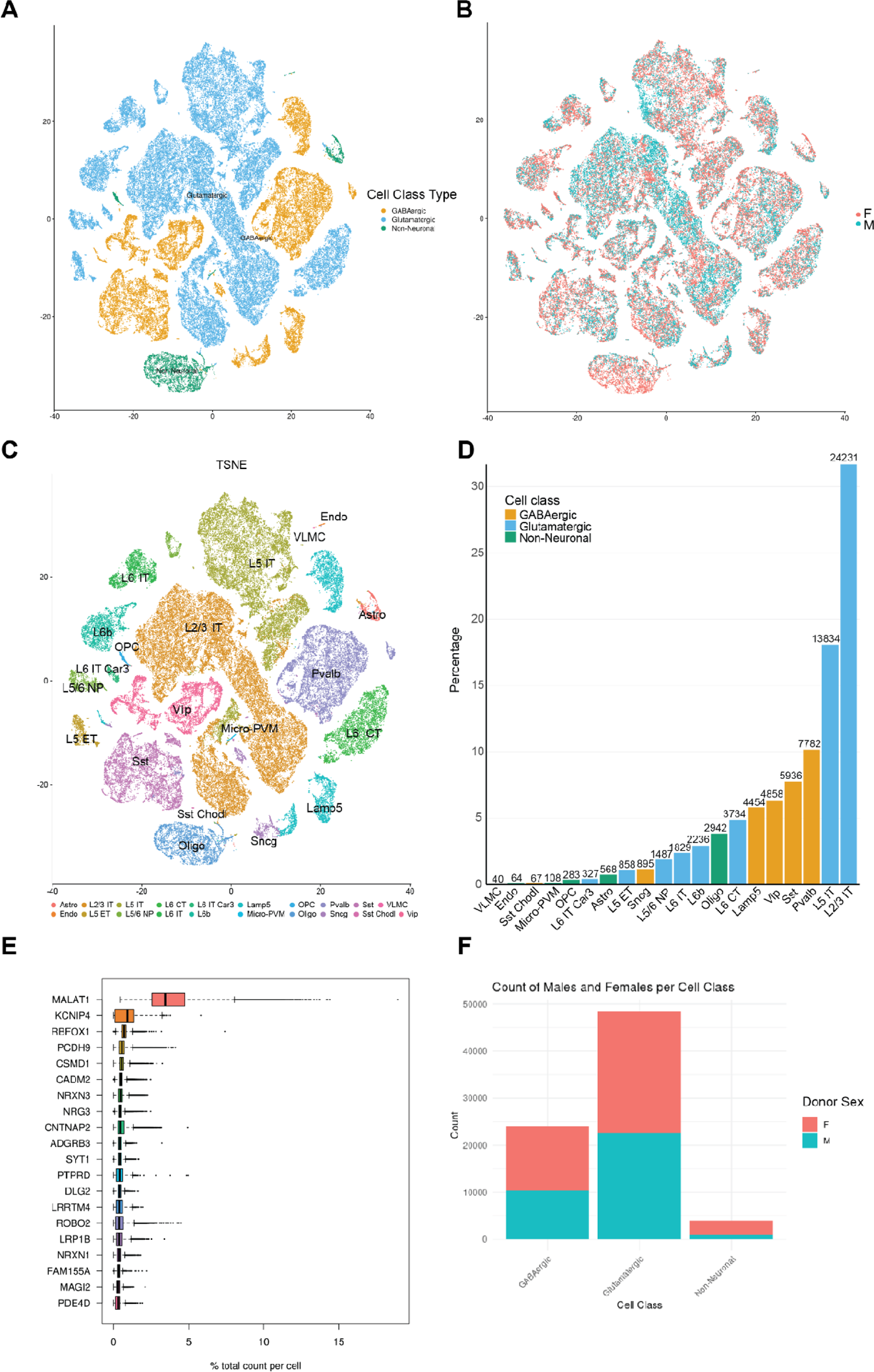
Single Cell Analysis Metrics of M1 Primary Motor Cortex from Single Cell Brain Atlas. **A.** t-SNE plot showing GABAergic, Glutamatergic and non-neuronal cell labels. **B.** t-SNE plot showing cells colored by gender. **C.** tSNE plot showing cell types found in the human brain atlas (Bakken et al. 2021). **D.** Bar plot shows the percentage of total cells and the count above each bar for each cell type. Bars are colored with cell class. **E.** Box plot sorted by the percentage of genes count expression per cell. **F.** Bar plot representing the differences in counts of cells by donor sex per cell class.

**Supplementary Figure 7:**
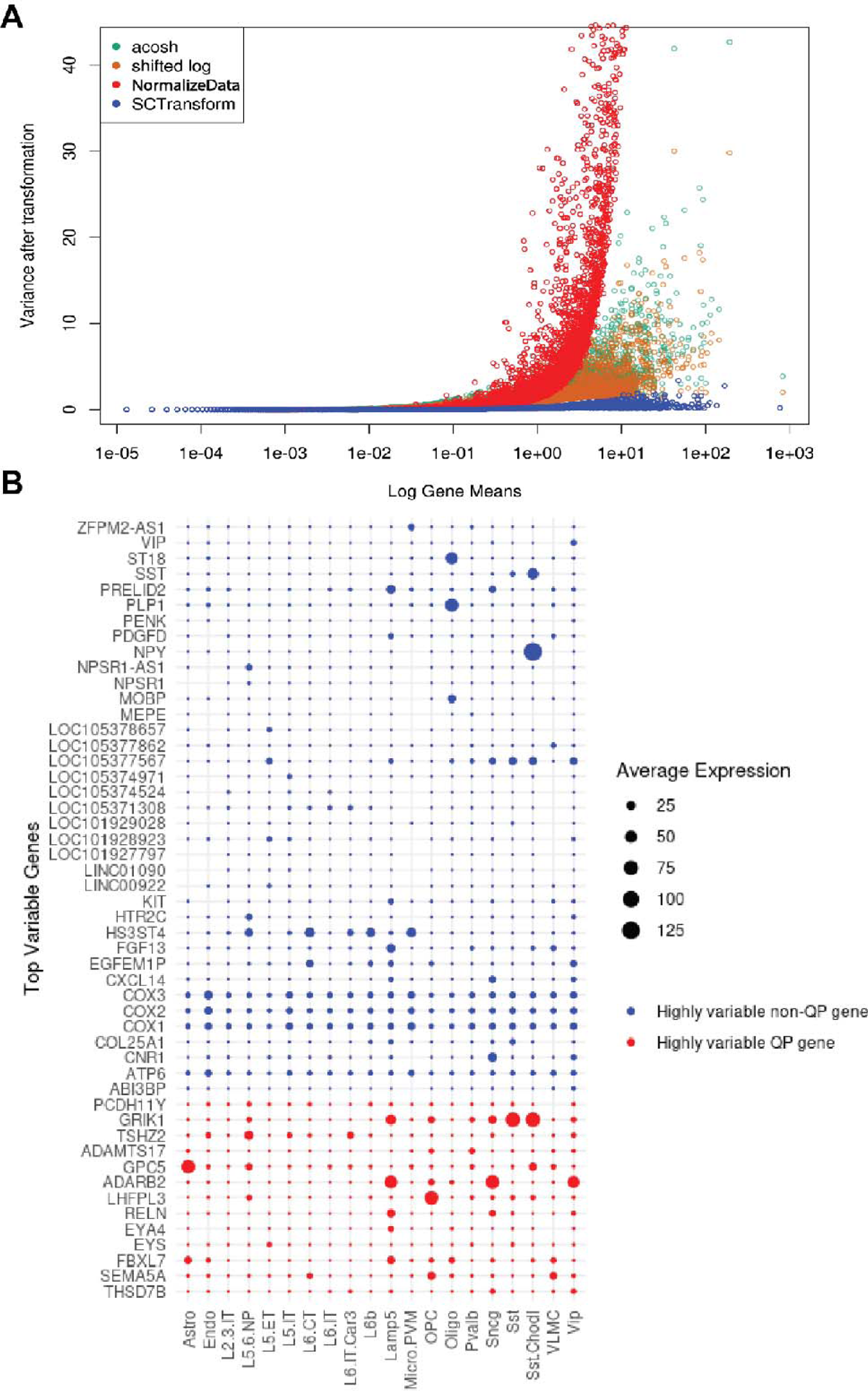
Single Cell Transformation and Variance Analysis. **A.** Variance after transformation using four methods, acosh, shifted log, NormalizeData (Seurat), and SCTransform (Seurat). **B.** Dot plot representing the average expression across cell types between quasi-prime and non-quasi prime genes for the top 25 genes in the dataset.

**Supplementary Figure 8:**
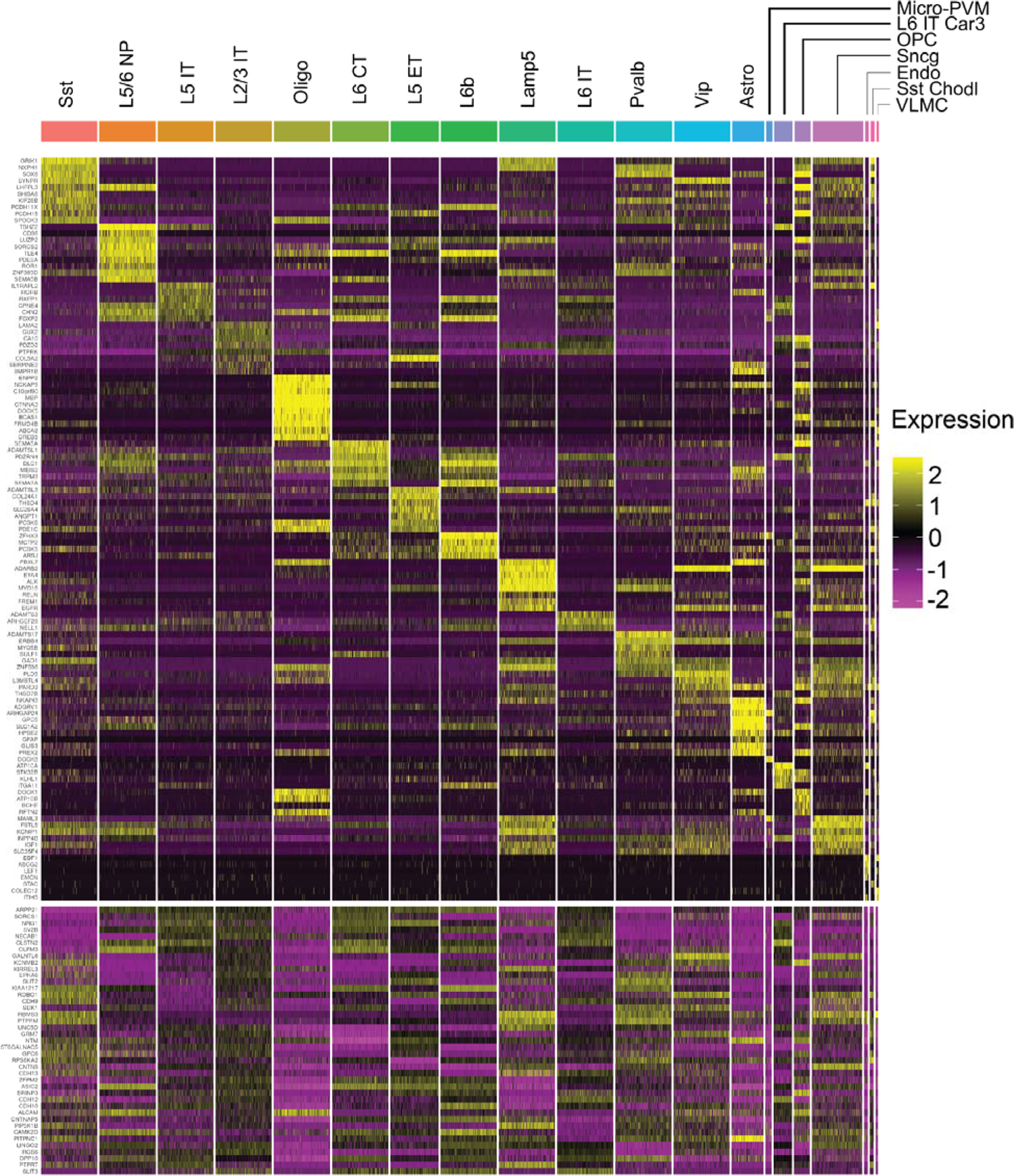
Differential Expression based on quasi-prime genes among cell types. **A.** Differentially expressed quasi-prime genes (p value < 0.05) were sorted by log2 fold change. The top 10 upregulated genes (log2 fold change > 0) and the top 10 downregulated genes (log 2 fold change < 0) from each cell type were selected and visualized in this heatmap. Upregulated and downregulated genes are separated in heatmap by row space.

**Supplementary Figure 9:**
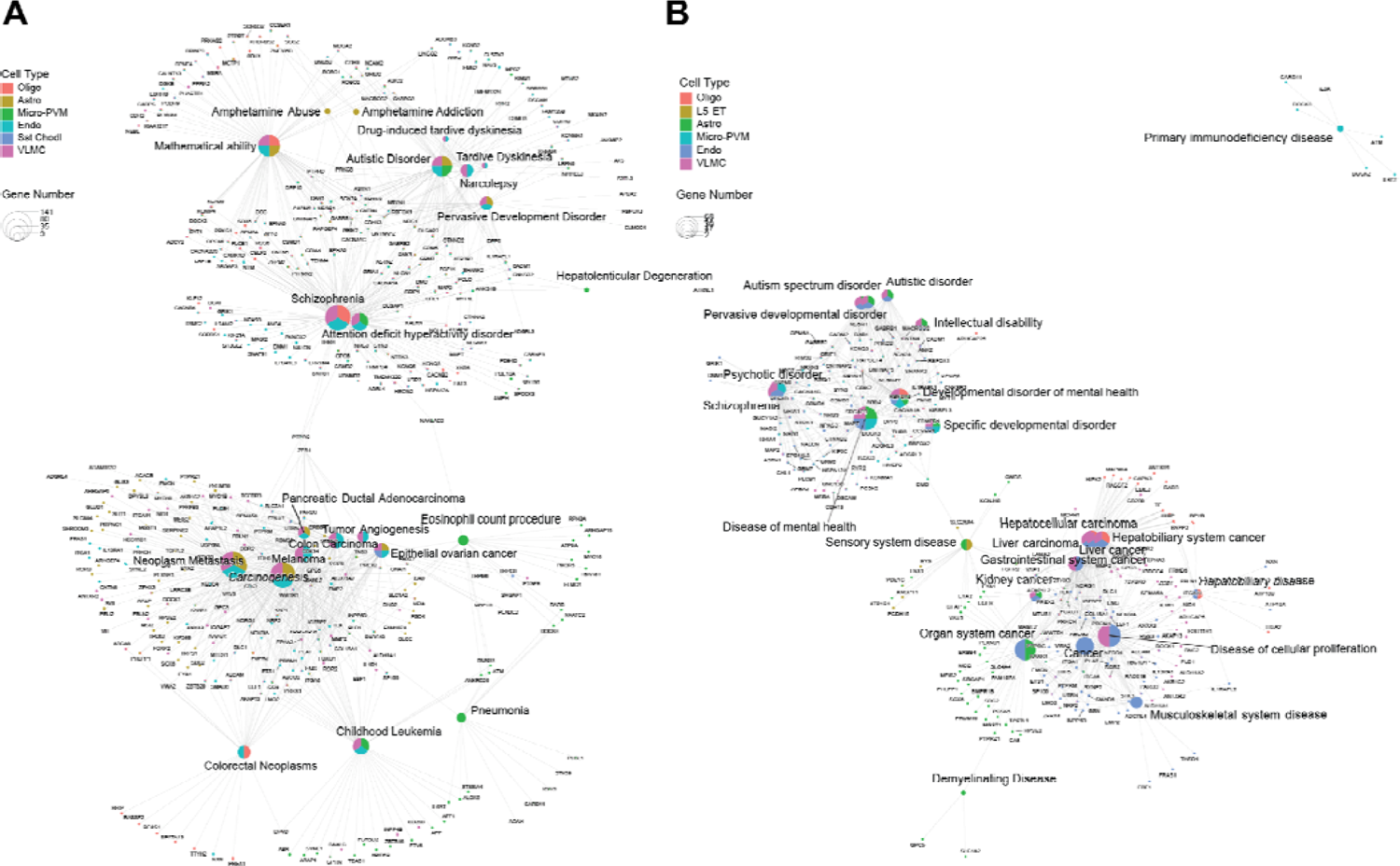
Gene set enrichment analysis of differentially expressed genes associated with quasi-prime sequences display several disease associations among cell types found in the human single-cell primary motor cortex brain atlas. **Significant (p value** < 0.05) differentially expressed genes with an absolute log2 fold change > 1 amongst cell types were used in GSEA analysis as represented by a network graph. **A.** DisGeNET “ALL” database. **B.** Disease ontology database. Different cell types are color coded and pie charts reflect the cell types at which each the diseases shown were associated.

**Supplementary Table 1:** UMLS Keywords and CUIs of interest used in disease2gene DisGeNET query.

| **Keywords** |  |
| --- | --- |
| disease\|disorder\|cancer\|neoplasm\|mental\|genetic\|drug\|neurologic\|brain\|health\|cognitive\|intelligence\|addiction\|dementia\|disability\|hereditary\|development\|deficiency\|condition\|itis\|infection\|ability\|microbiome\|microbiota\|pneumonia\|cirrhosis\|fibrosis\|acquired\|resistant\|syndrome\|degeneration\|pathology\|pathologic\|behavior\|psych\|dysfunction\|abnormal\|virus\|bacteria\|motor\|paralysis\|defect\|human |  |
| **Disease Class** | **CUIs** |
| Psychosis/Mental/Cognitive/Nervous System Disorders and Brain Dysfunction | "C0036341", "C0002395", "C0005586", "C3714756", "C0014544", "C0036572", "C1510586", "C0004352", "C0011581", "C0011570", "C0027765", "C0751872", "C0004930", "C0011570" |
| Substance and Drug Abuse | "C0013146", "C0001973" |
| Degenerative and Autoimmune Disorders | "C0002736", "C0030567", "C0003873", "C0021053", "C0014130", "C0029456" |
| Neurodevelopmental Disorder | "C1535926", "C0557874" |
| Neoplastic Processes | "C0023467", "C0026998", "C0023418", "C0006826" |
| Lung Diseases | "C0024115" |
| Eye Disorders | "C0015397", "C0242383" |
| Metabolic Diseases | "C0011849", "C0020538" |
| Vascular Diseases | "C0007222", "C0007820" |
| Additional Human Specific Diseases | "C0019163", "C0021400", "C0025007", "C0043167", "C0035869", "C0039128", "C0039614", "C0041296", "C0001175", "C0041234", "C0008354", "C0019104", "C0011311", "C0041228", "C0024537", "C0024535", "C0024530", "C0024534", "C0023290", "C1301422", "C0026780", "C0032064", "C0035920", "C0037354", "C0041466", "C0041471", "C0043395", "C0035235", "C0035473", "C0001487", "C0010823", "C0155862", "C0032371", "C0010246", "C0019348", "C0036439", "C0410702", "C1837461" |

## Notes

### Competing Interest Statement

The authors have declared no competing interest.

